# A model-free method for measuring dimerization free energies of CLC-ec1 in lipid bilayers

**DOI:** 10.1101/154286

**Authors:** Rahul Chadda, Lucy Cliff, Marley Brimberry, Janice L. Robertson

**Affiliations:** Department of Molecular Physiology and Biophysics, The University of Iowa, Iowa City IA.; Department of Chemistry, The University of Bath, Bath, UK.

## Abstract

We previously reported the equilibrium dimerization reaction of the CLC-ec1 Cl^-^/H^+^ transporter in 2:1 POPE/POPG membranes (Chadda et al. 2016). This was determined by measuring the probability distributions of subunit capture into extruded liposomes by single-molecule photobleaching analysis across a wide range of subunit/lipid mole fraction densities. In this approach, knowledge of the liposome size distribution is necessary in order to correct the data for random co-capture events and extract the underlying dimerization reaction. For this we used a previously reported cryo-electron microscopy (cryo-EM) measured size distribution of 400 nm extruded liposomes made of *E. coli* polar lipids (Walden et al. 2007). While the model and data agreed at low densities, we observed systematic inaccuracies at higher densities limiting our ability to extract *F_Dimer_* in this range. To address this issue, we measured the 400 nm extruded 2:1 POPE/POPG liposome size distribution by cryo-EM and found that there is a small, but significant amount of larger liposomes in the population. Re-analysis of the I201W/I422W ‘WW’ photobleaching data using this distribution shows that the protein is monomeric in the membrane and can serve as an experimental control. Dimer controls were constructed by glutaraldehyde cross-linking of C85A/H234C ‘WT’ or introducing R230C/L249C, which forms a spontaneous disulfide bond. Determination of *F_Dimer_* based on the experimental controls yields improved fits and no change in the previously reported *ΔG*° values, providing an alternate model-free approach to measuring CLC-ec1 dimerization in membranes.

## RESULTS

Equilibrium association reactions of stable membrane protein complexes in lipid bilayers have been challenging to measure due to limited protein signals. This has restricted studies to that of weak complexes (Yano & Matsuzaki 2006; Cristian et al. 2011; North et al. 2006; Yano et al. 2015), though advances into steric-trapping have enabled the study of stable oligomers (Hong et al. 2010; Hong & Bowie 2011). For an alternate approach, it has been demonstrated that reconstitution of membrane proteins into liposomes can report on protein stoichiometry at low densities in the membrane (Walden et al. 2007; Robertson et al. 2010; Stockbridge et al. 2013; Fang et al. 2006). Using this approach and taking advantage of the sensitivity of single-molecule photobleaching analysis, we were able to measure CLC-ec1 stoichiometry at various subunit/lipid densities, including extremely dilute conditions where we observed a shift from dimer to monomer (Chadda et al. 2016; Chadda & Robertson 2016). The fraction of protein in the dimer state showed a reversible dependency on the membrane density, allowing for the determination of the dimerization free energy in lipid bilayers.

The major challenge in this approach is to measure population stoichiometry accurately across a wide range of protein densities. In order to observe a complete change in the population from monomers to dimers, we must examine the stoichiometry across five orders of magnitude, and at densities greater than the Poisson limit where liposomes are likely to contain more than one protein species. This issue can be exacerbated in a heterogenous liposome population as larger liposomes tend to act like protein sinks. This random co-capture of protein subunits has been demonstrated to falsely report on oligomerization for membrane proteins in detergent micelles (Tanford & Reynolds 1976; Stanley & Fleming 2005), and it must be corrected for in any reconstituted system. Previously, we used a stochastic simulation of the Poisson process of subunit reconstitution into a defined liposome population based on the ‘Walden’ distribution of 400 nm extruded vesicles comprised of *E. coli* polar lipids (EPL) (Figure 1A,B) (Walden et al. 2007). While the data and model agreed at lower densities, it systematically deviated at higher densities where the data contained fewer single-steps and more multi-step photobleaching events as compared to the model (Figure 1C,D). We hypothesized that the larger liposomes were under-represented in the Walden distribution leading to an underestimation of liposomes containing more than 3 steps in the model. We directly investigated this by measuring the 400 nm extruded 2:1 POPE/POPG liposome size distribution by cryo-electron microscopy (Figure 1E), which shows that there is a significant proportion of larger liposomes that is missing in the Walden distribution. The difference is small but the effect is amplified when considering the fractional surface area (Figure 1F), which dictates the Poisson process of subunit capture. This underrepresentation of larger liposomes in the Walden distribution could arise due to size selection during the freezing of cryo-electron microscopy samples, or it could be a result of minor differences in the lipid composition. EPL is a crude extract with approximately 67% POPE, 20% POPG and 10% cardiolipin, while our experimental lipid conditions is a synthetic mimic made of 67% POPE and 23% POPG.

**Figure 1.**
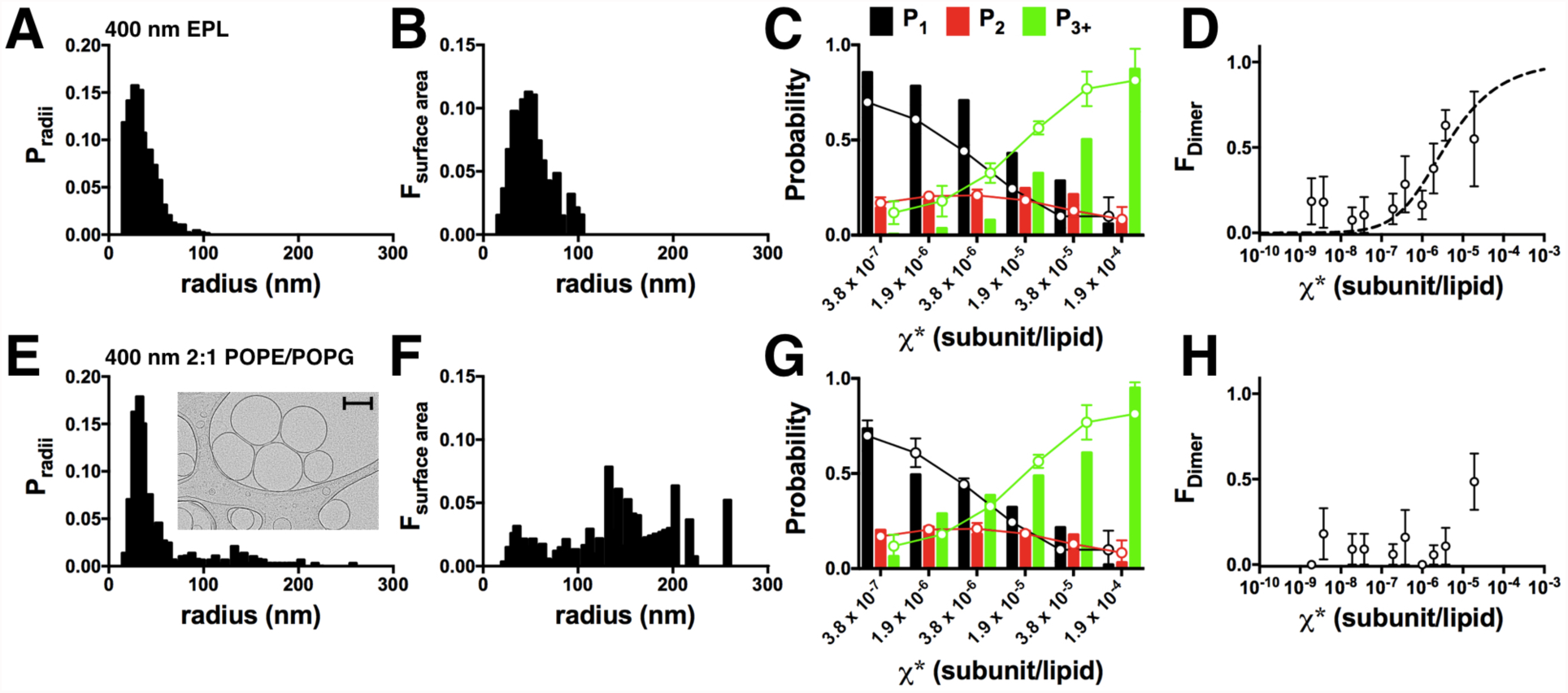
The 400 nm 2:1 POPE/POPG liposome size distribution indicates WW-Cy5 is monomeric. (A) 400 nm EPL liposome size distribution reported in Walden et al. (Walden et al., 2007). (B) Fractional surface area distribution. (C) Photobleaching distribution of model (bars) and experimental WW data (circles-lines). (D) F_Dimer_ values for WW-Cy5 determined using the Walden distribution, reported in Chadda et al. (Chadda et al., 2016). (E) The 400 nm 2:1 POPE/POPG liposome size distribution (mean, n=2). Inset shows a cryo-electron microscopy image of the liposomes (raw image attached as source data), scale bar - 200 nm. (F) Fractional surface area distribution (F_surface area_). (G) Photobleaching distribution model (bars) and experimental WW data (circles-lines). (H) Re-calculated F_Dimer_ values for WW-Cy5 from the updated 2:1 POPE/POPG distribution, demonstrating that WW-Cy5 is monomeric across the dynamic range of the experiment. Data represented as mean ± SEM, n=2-3.

With the updated liposome size distribution, we re-examined the I201W/I422W (‘WW’) CLC-ec1 photobleaching data from Chadda et al. (Chadda et al. 2016). Previously, this construct was found to be monomeric in detergent by both glutaraldehyde cross-linking, x-ray crystallography and also in 3:1 egg PC/POPG liposomes reconstituted at *χ* = 1.5 x 10^−5^ subunits/lipid (1 μg/mg) (Robertson et al. 2010). We also observed differences in the fraction of empty liposomes (F_0_) by single-molecule co-localization microscopy for the Cy5 labeled protein and Alexa Fluor 488 labeled liposomes, indicating that the protein occupancy was consistent with a monomer at high densities (Chadda et al. 2016). However, when we calculated *F_Dimer_* using the Walden distribution, we observed a weak apparent dimerization reaction that either indicated an actual formation of dimers or inaccuracies of the modeling at higher densities. With the new liposome distribution, we find that the experimental data converges with the simulated monomer probabilities (Figure 1G) and that the apparent dimerization is no longer present (Figure 1H). This, together with the other evidence presented in previous studies, demonstrates that WW is monomeric within our experimental range of densities. This observation indicates that WW can serve as a monomeric control in future experiments.

With a monomeric control in place, we were motivated to identify an experimental dimer control, to establish a model-free approach to measuring dimerization that does not require prior knowledge of the liposome size distribution. For this, we turned to covalent cross-linking methods that have already been well established for CLC-ec1. First, we tried glutaraldehyde that covalently cross-links the dimer state as demonstrated on SDS-PAGE (Figure 2 – figure supplement 1) (Maduke et al. 1999; Robertson et al. 2010). Glutaraldehyde is a short chain bis-reactive molecule and cross-links proteins via primary amine groups present on lysines and the N-terminus. CLC-ec1 has 13 native lysine residues (Figure 2A) and while the dimer is the major cross-linked product, we also observe a small tetramer population and a small amount of resistant monomer (Figure 2 - supplementary figure 1), highlighting the non-specific nature of this reaction. Measurement of the photobleaching probability distribution shows that the WT-Cy5 + glutaraldehyde proteoliposomes follow the 2:1 POPE/POPG dimer model, as well as the saturating range of the WT-Cy5 photobleaching data (Figure 2B). However, we observe lower values of P_1_ and higher values of P_3+_, which agrees with the observation that a small fraction of protein is non-specifically cross-linked in a higher oligomeric state. This contribution, however, is small and the majority of the reconstituted protein is dimeric. Upon measurement of functional activity, however, it was found that a large fraction of the protein is non-functional (Figure 2 – figure supplement 1). Therefore, while the protein is present in the membrane, in mainly a dimeric form, the background glutaraldehyde modifications or cross-linking states have a significant effect on transport function. With this in mind, we reserve glutaraldehyde cross-linked WT as a structural dimeric control in the membrane, but not one with a proper biological fold.

**Figure 2.**
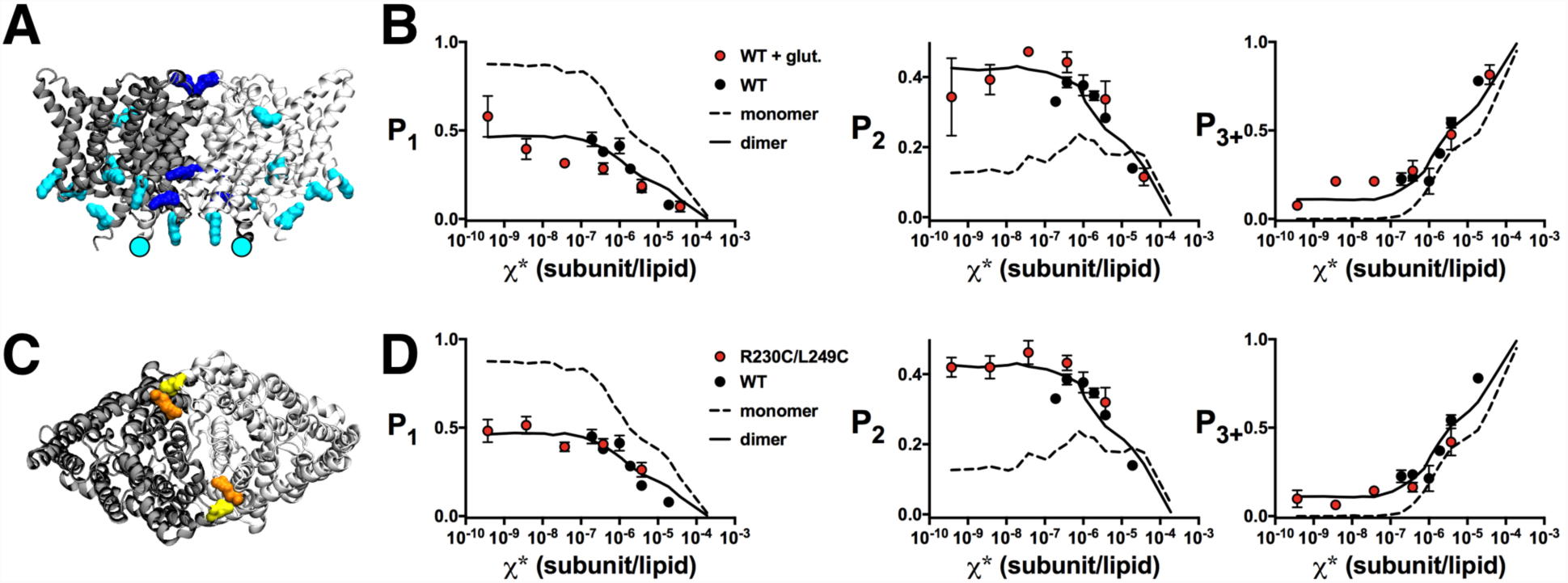
Photobleaching analysis of cross-linked CLC dimers. (A) Side view of CLC-ec1 homodimer, with the two subunits colored light and dark grey. Residues with primary amine groups that are potential glutaraldehyde cross-linking sites are highlighted. Lysines are turquoise with those within 5 Å of another lysine colored dark blue. The N-terminus is highlighted as a circle. (B) Photobleaching data of WT-Cy5 + glutaraldehyde (red) compared to the saturating range of WT-Cy5 (black) and monomer/dimer models using the 2:1 POPE/POPG liposome size distribution. Data are represented as mean ± SEM, n = 4–6, P_Cy5_ = 0.70 ± 0.04. (C) Top view of the CLC-ec1 homodimer showing R230C (orange) and L249C (yellow) positions. (D) Photobleaching data of R230C/L249C-Cy5 (red) compared to the saturating range of WT-Cy5 (black) and monomer/dimer models using the 2:1 POPE/POPG liposome size distribution. Data are represented as mean ± SEM, n=5, P_Cy5_ = 0.72 ± 0.02. Model parameters set as: P_Cy5_ = 0.72, P_non-specific_ = 0.14, bias = 4 and A_lipid_=0.6.

For an alternate approach, we investigated disulfide cross-linking across the dimerization interface. Previously, Nguitragool & Miller demonstrated that the CLC-ec1 dimer could spontaneously cross-link via a disulfide bond between R230C and L249C, during expression and/or purification (Nguitragool & Miller 2007). We introduced R230C/L249C onto the C85A/H234C ‘WT’ background (Figure 2C) and found that the protein expressed, purified as a dimer in detergent micelles (Figure 2 – figure supplement 2), and ran as a dimer on SDS-PAGE indicating disulfide formation prior to purification (Figure 2C). The disulfide bond is not modified by the reducing agent tris(2-carboxyethyl)phosphine (TCEP) included in the purification, which allows for H234C to remain reactive for Cy5-maleimide labeling at yields comparable to the WT. The photobleaching probability distributions were measured for 2:1 POPE/POPG membranes incubated at room temperature for 1-4 days and then extruded using a 400 nm filter (Figure 2D). The data shows that the photobleaching probability distribution of R230C/L249C-Cy5 corresponds to the ideal dimer simulation based on the updated 2:1 POPE/POPG liposome size distribution, as well as the saturating range of the WT-Cy5 data. In addition, we measured the chloride transport function of the R230C/L249C proteoliposomes reconstituted at *χ* = 1.5 x 10^−5^ subunits/lipid, which showed comparable function to WT (Figure 2 – figure supplement 2). Therefore, R230C/L249C provides a functionally competent dimer control for CLC-ec1 dimerization reactions, preserving the native functional fold.

With these non-reactive, ‘ideal’ monomer and dimer experimental controls, we re-calculated *F_Dimer_* for the WT-Cy5 and W-Cy5 photobleaching data. The fits of the equilibrium dimerization isotherm (Figure 3) are improved using either WT + glutaraldehyde or R230C/L249C data for the dimer state and WW data defining the monomeric state. The values of *ΔG*° and *ΔΔG* are not significantly different compared to our previous report (Figure 3E), showing that the simulation method is sufficient to estimate *F_Dimer_* but that the physical controls provide a model-free method of obtaining the same results.

**Figure 3.**
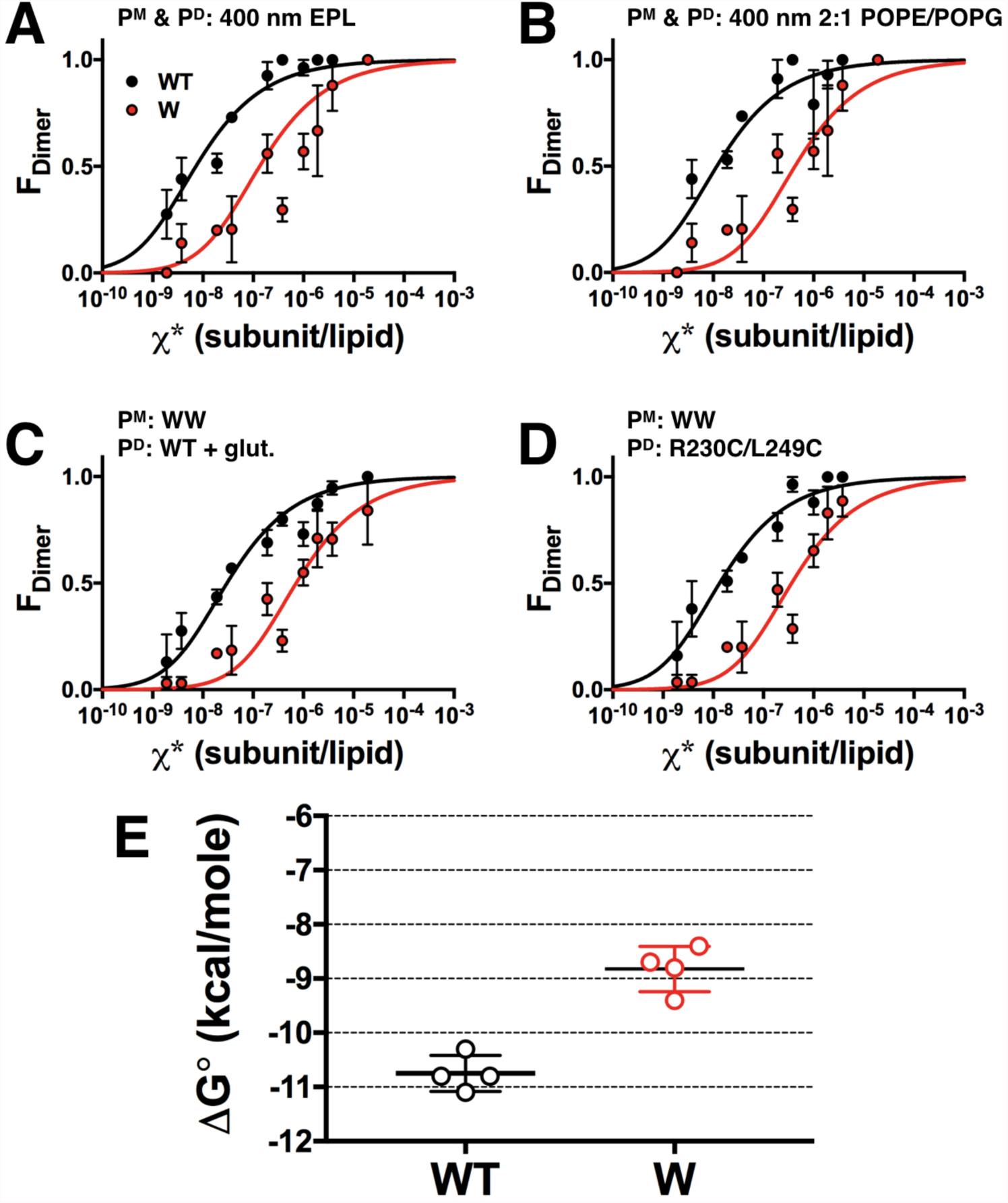
Model-free measurements of WT-Cy5 and W-Cy5 dimerization free energies in 2:1 POPE/POPG lipid bilayers. F_Dimer_ analysis of WT-Cy5 (black) and W-Cy5 (red). Monomer and dimer probabilities determined by (A) modeling using the 400 nm EPL Walden liposome size distribution (model parameters set as: P_Cy5_ = 0. 72, P_non-specific_ = 0. 14, bias = 4 and A_lipid_=0.6), (B) modeling the 400 nm 2:1 POPE/POPG liposome size distribution, (C) WW-Cy5 for monomer and glutaraldehyde cross-linked WT-Cy5 for dimer, (D) WW-Cy5 for monomer and R230C/L249C-Cy5 for dimer. Data represent mean ± SEM, n=2-3. (E) Comparison of ΔG° values for all monomer/dimer models and controls, mean ± SD. Values are reported in Table 1, and figure supplements 1 & 2.

**Table 1.** Summary of free energy values for CLC-ec1 dimerization in 2:1 POPE/POPG lipid bilayers. Free energies are calculated as ΔG° = -RT ln(K_eq_/χ°) where χ° is the standard state of 1 subunit/lipid, R is the gas constant 1.99 x 10^−3^ kcal mole^-1^ K^-1^, and T is 298 K (25 °C), where K_eq_ is determined by fitting to the equilibrium dimerization isotherm. Data represent bestfit ± error of fit (R^2^).

| Monomer/dimer data | $\Delta G^\circ_{WT}$ (kcal/mole) | $\Delta G^\circ_W$ (kcal/mole) |
| --- | --- | --- |
| P<sup>M</sup>: 400 nm EPL, $K_D = 1 \times 10^{100}$<br>P<sup>D</sup>: 400 nm EPL, $K_D = 1 \times 10^{-100}$ | $-11.1 \pm 0.1$ (0.91) | $-9.4 \pm 0.2$ (0.87) |
| P<sup>M</sup>: 400 nm 2:1 POPE/POPG,<br>$K_D = 1 \times 10^{100}$<br>P<sup>D</sup>: 400 nm 2:1 POPE/POPG,<br>$K_D = 1 \times 10^{-100}$ | $-10.8 \pm 0.2$ (0.80) | $-8.7 \pm 0.2$ (0.71) |
| P<sup>M</sup>: WW<br>P<sup>D</sup>: WT + glutaraldehyde | $-10.3 \pm 0.1$ (0.88) | $-8.4 \pm 0.2$ (0.78) |
| P<sup>M</sup>: WW<br>P<sup>D</sup>: R230C/L249C | $-10.8 \pm 0.1$ (0.89) | $-8.8 \pm 0.2$ (0.80) |

This study reports on methodological advances developed for measuring dimerization of CLC-ec1 in 2:1 POPE/POPG lipid bilayers. Using a newly determined liposome size distribution, we increase the dynamic range of the measurement one order of magnitude up to *χ* = 3.8 x 10^−5^ subunits/lipid, allowing for stability measurements of weaker complexes. We also establish experimental monomeric (WW) and dimeric (WT + glutaraldehyde or R230C/L249C) controls that provide an easy way of correcting the experimental data for random co-capture without knowledge of the liposome size distribution. This simplifies measurement of the reaction under changing conditions, for instance temperature and lipid composition, both of which are expected to change the liposome size distribution. The overall agreement between the statistical modeling approach and experimental controls demonstrates that the subunit capture and accounting approach is sufficiently rigorous for reporting on membrane protein stoichiometry in lipid bilayers.

## METHODS

The bulk of the methods used in this study follow that reported in (Chadda et al. 2016). Details of experiments specific to this study are outlined here.

### Cross-linking of ‘WT’ C85A/H234C CLC-ec1

For running on SDS-PAGE, glutaraldehyde (Sigma Aldrich, St. Louis, MO) was added to 8 μM WT in size exclusion buffer (SEB; 150 mM NaCl, 20 mM MOPS pH 7.5, 5 mM analytical-grade DM, Anatrace, Maumee, IL), for a final concentration of glutaraldehyde of 0.4% wt/vol (~40 mM). The reaction was allowed to proceed for 8 minutes after which 10x Tris or Glycine buffer was added to quench the reaction. For reconstitution, ‘WT’ protein was labeled with Cy5-maleimide as described previously, then cross-linked with glutaraldehyde and quenched as described above, before reconstitution into 2:1 POPE/POPG liposomes (Avanti Polar Lipids, Alabaster, AL). For the R230C/L249C disulfide cross-linked construct (Nguitragool & Miller 2007), mutations were added to the C85A/H234C background using a Quickchange II site-directed mutatgenesis kit (Agilent Technologies, Santa Clara, CA). Purification was carried out as described previously (Chadda et al. 2016), in the presence of 1 mM TCEP until the SEC purification step. Labeling and reconstitution was carried out as before. For SDS-PAGE, the sample was run on non-reducing gels. For DTT reduction, 10 μM of protein was incubated with 100 mM DTT at 30 °C for 1 hour.

### Cryo-electron microscopy measurements of liposome size distributions

Liposomes were freeze/thawed seven times, incubated at room temperature, and then extruded through a 400 nm nucleopore filter (GE Life Sciences, Chicago, IL) 21 times prior to sample freezing. For freezing, 3 μL of the undiluted sample was loaded onto glow-discharged Lacey carbon support films (Electron Microscope Sciences, Hatfield, PA), blotted and plunged into liquid ethane using a Vitrobot System (FEI, Hilsboro, OR). Images were collected at 300 kV on a JEOL 3200 fs microscope (JEOL, Tokyo, Japan) with a Gatan K2 Summit direct electron detector camera (GATAN, Pleasanton, CA). Magnifications of 15K and 30K were used. Liposome sizes were analyzed in Fiji & ImageJ (Schneider et al. 2012; Schindelin et al. 2012), and all liposomes were included in counting, even those on the carbon outside of the vitreous ice.

### *F_Dimer_* calculator

A MATLAB app was created to calculate the fraction of dimer using the various models and experimental controls, and the sum of *R^2^* analysis. All of the models and experimental control data are implemented in the code. For the modeling, *P_Cy5_*= 0.72, *P_non-specific_*= 0.14 was used, and a bias 4 for size exclusion of dimer model, A_lipid_ = 0.6 nm^2^ as described in (Chadda et al. 2016; Chadda & Robertson 2016). The app file is available for download as a source file in the supplementary information. MATLAB 2016b or higher is required.

## ACKNOWLEDGEMENTS

We acknowledge Thomas Moninger at the University of Iowa Microscopy Core Facility, and Jonathan Remis, staff and instrumentation support at the Structural Biology Facility at Northwestern University. The Structural Biology Facility is partially supported by the R.H. Lurie Comprehensive Cancer Center of Northwestern University.

---

**Figure 1 - figure supplement 1.**
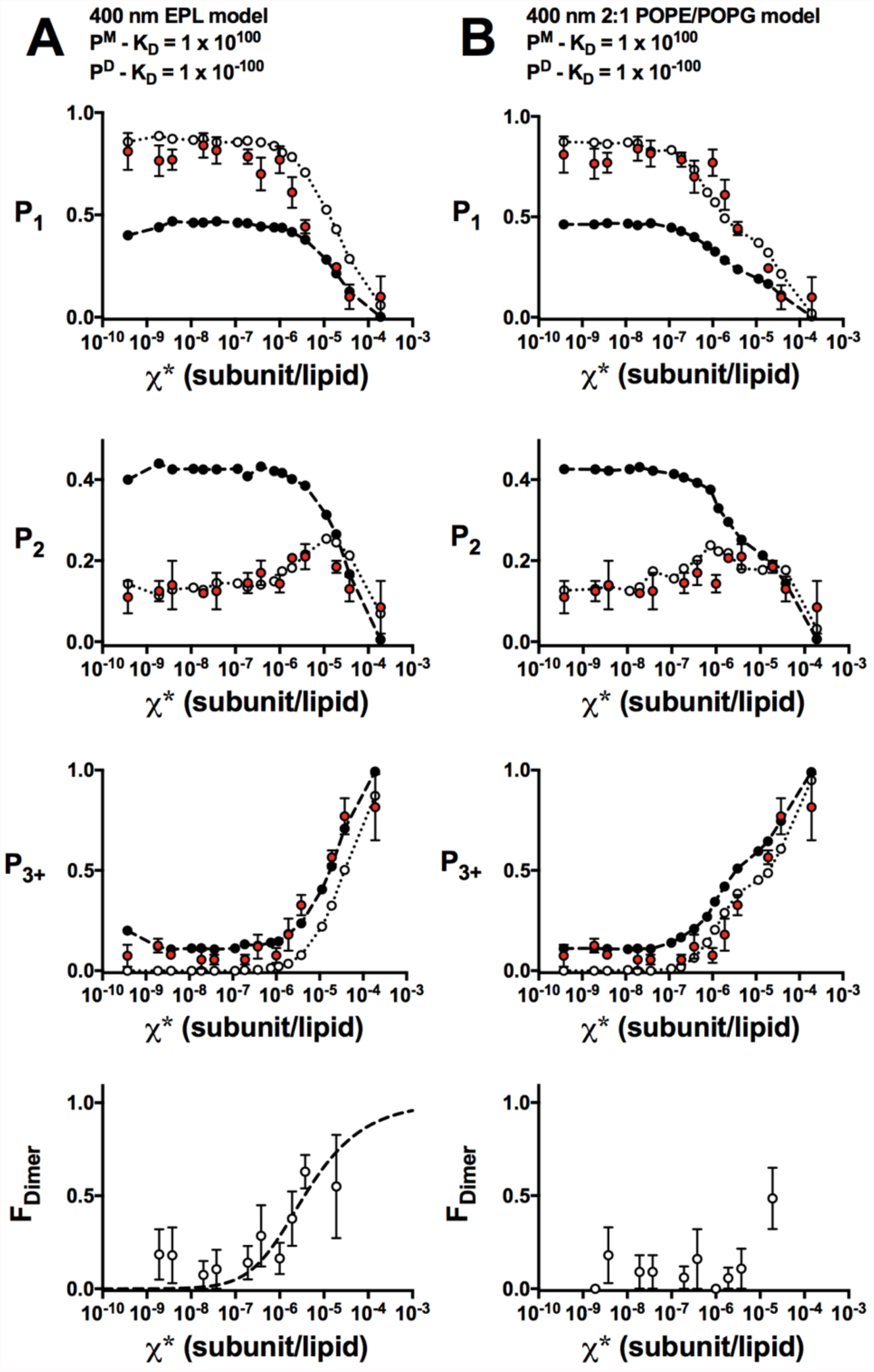
F_Dimer_ analysis of WW-Cy5 in 2:1 POPE/POPG membranes. Photobleaching probabilities (P_1_, P_2_, P_3+_) and F_Dimer_ analysis (bottom row) of WW-Cy5 (red) along with monomer (white) and dimer (black) probabilities. Monomer and dimer probabilities determined by (A) modeling using the 400 nm EPL Walden liposome size distribution, ΔG° = -7.4 ± 0.2 kcal/mole, R^2^ = 0.30 (B) modeling using the 400 nm 2:1 POPE/POPG liposome size distribution, no fit shown. Model parameters set as: P_Cy5_ = 0.72, P_non-specific_ = 0. 14, bias = 4 and A_lipid_=0.6. Data represent mean ± SEM, n=2-3.

**Figure 2 - figure supplement 1.**
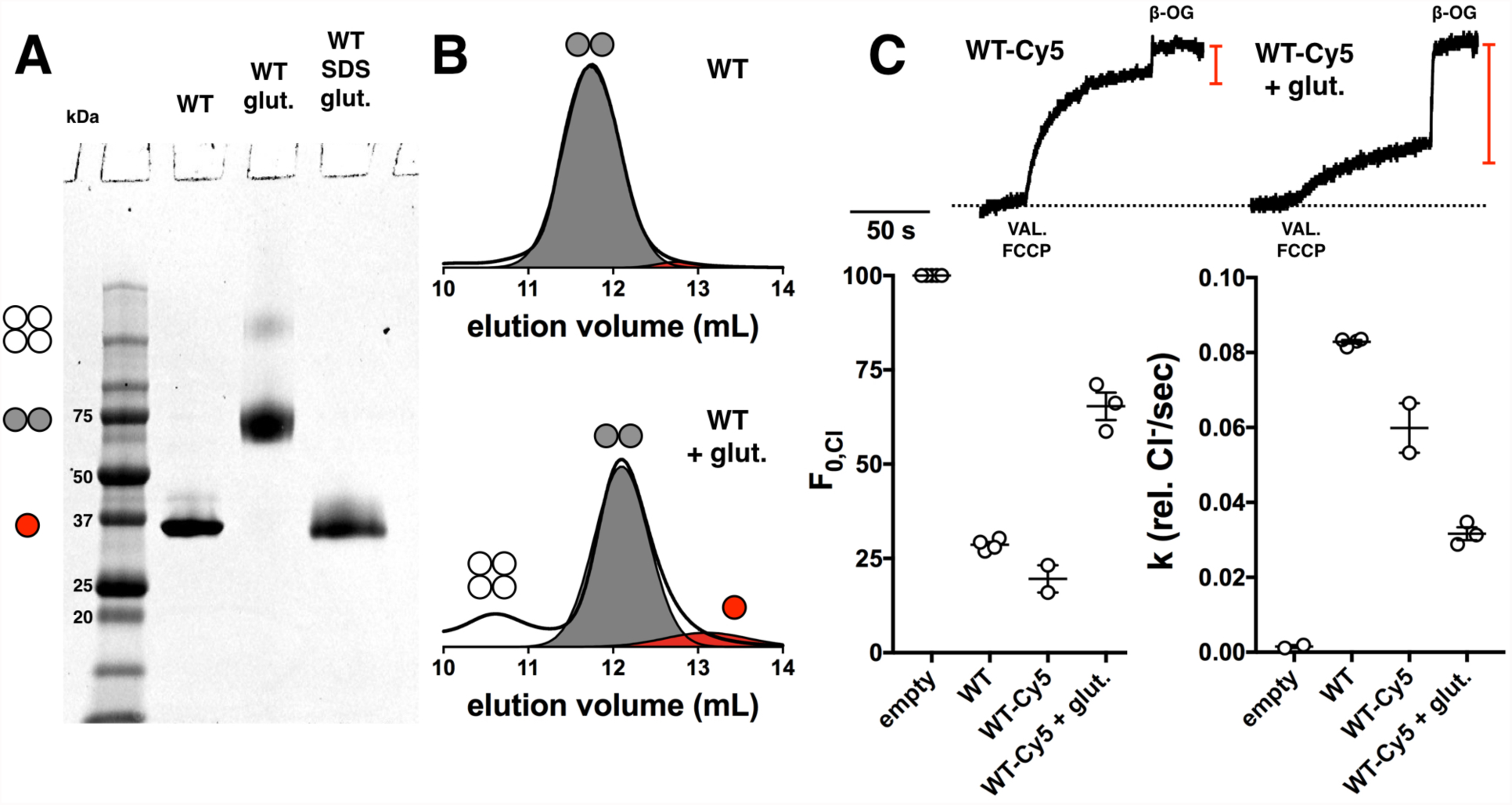
Glutaraldehyde cross-linking of WT-Cy5. (A) SDS-PAGE of WT, WT + glutaraldehyde, WT + 2% SDS followed by addition of glutaraldehyde. Red circle – monomer, grey circles – dimer, white circles - tetramer. (B) Size exclusion chromatography profiles of WT and glutaraldehyde cross-linked WT. (C) Functional Cl^-^ transport of WT-Cy5 and WT-Cy5 + glutaraldehyde, reconstituted at 1 μg/mg (χ = 1.5 x 10^−5^ subunits/lipid). Data represented as mean ± SEM. The fractional volume of empty liposomes (F_0,Cl,_ red bar) and transport rate (k) shows a significant reduction in CLC function in the presence of glutaraldehyde (*p*<0.0001 &*p*=0.009 respectively).

**Figure 2 - figure supplement 2.**
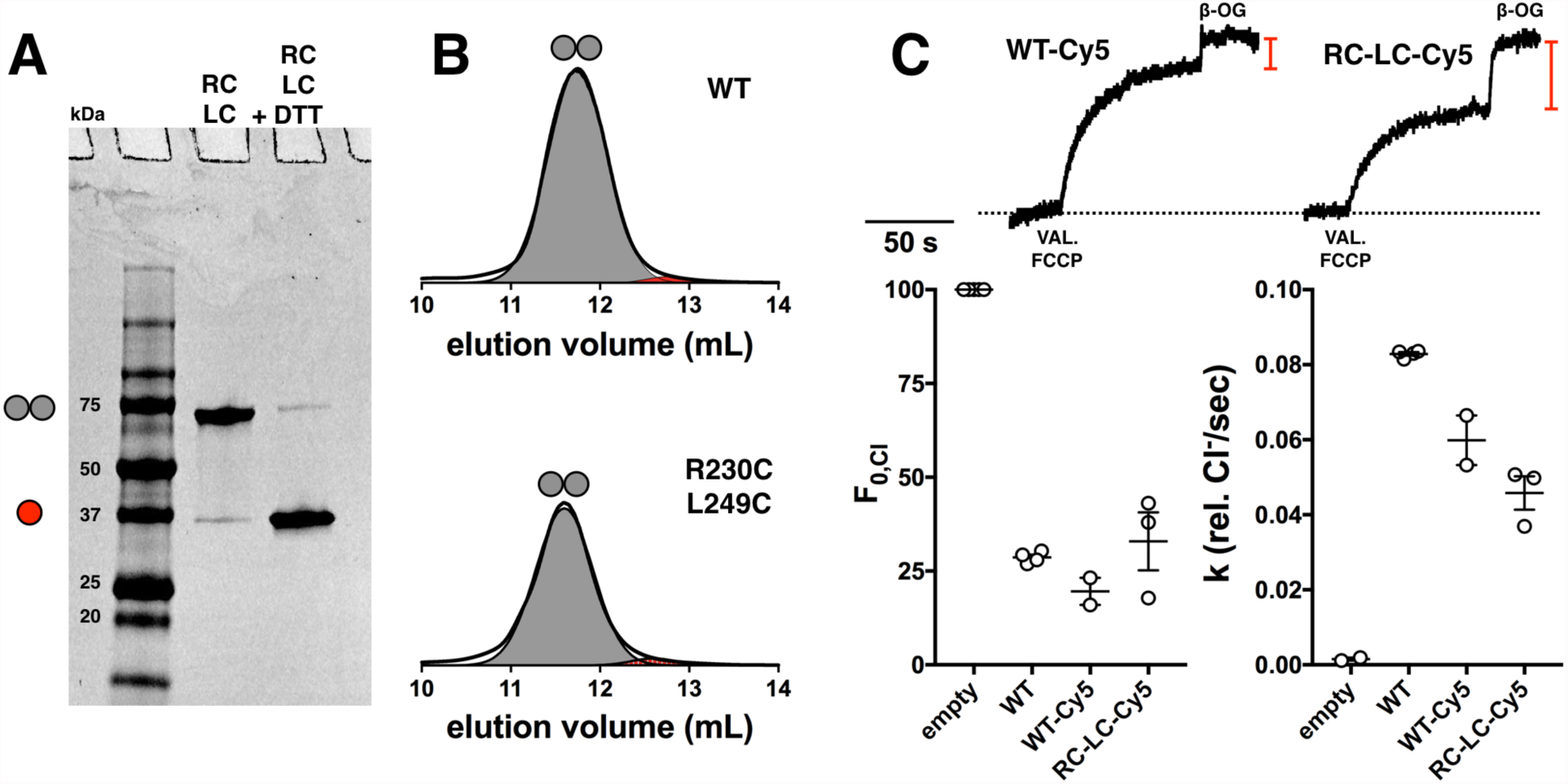
Disulfide cross-linking of CLC-ec1. (A) SDS-PAGE of R230C/L249C and R230C/L249C treated with DTT. Red circle – monomer, grey circles – dimer. (B) Size exclusion chromatography profiles of WT and R230C/L249C. (C) Functional Cl transport of WT-Cy5 and R230C/L249C-Cy5 (RC-LC-Cy5), reconstituted at 1 μg/mg, χ = 1.5 x 10^−5^ subunits/lipid. Data represented as mean ± SEM. The fractional volume of empty liposomes (F_0,Cl_, red bar) and Cl^-^ transport rate (k) show no significant change between WT-Cy5 and RCLC-Cy5 samples.

**Figure 3 - figure supplement 1.**
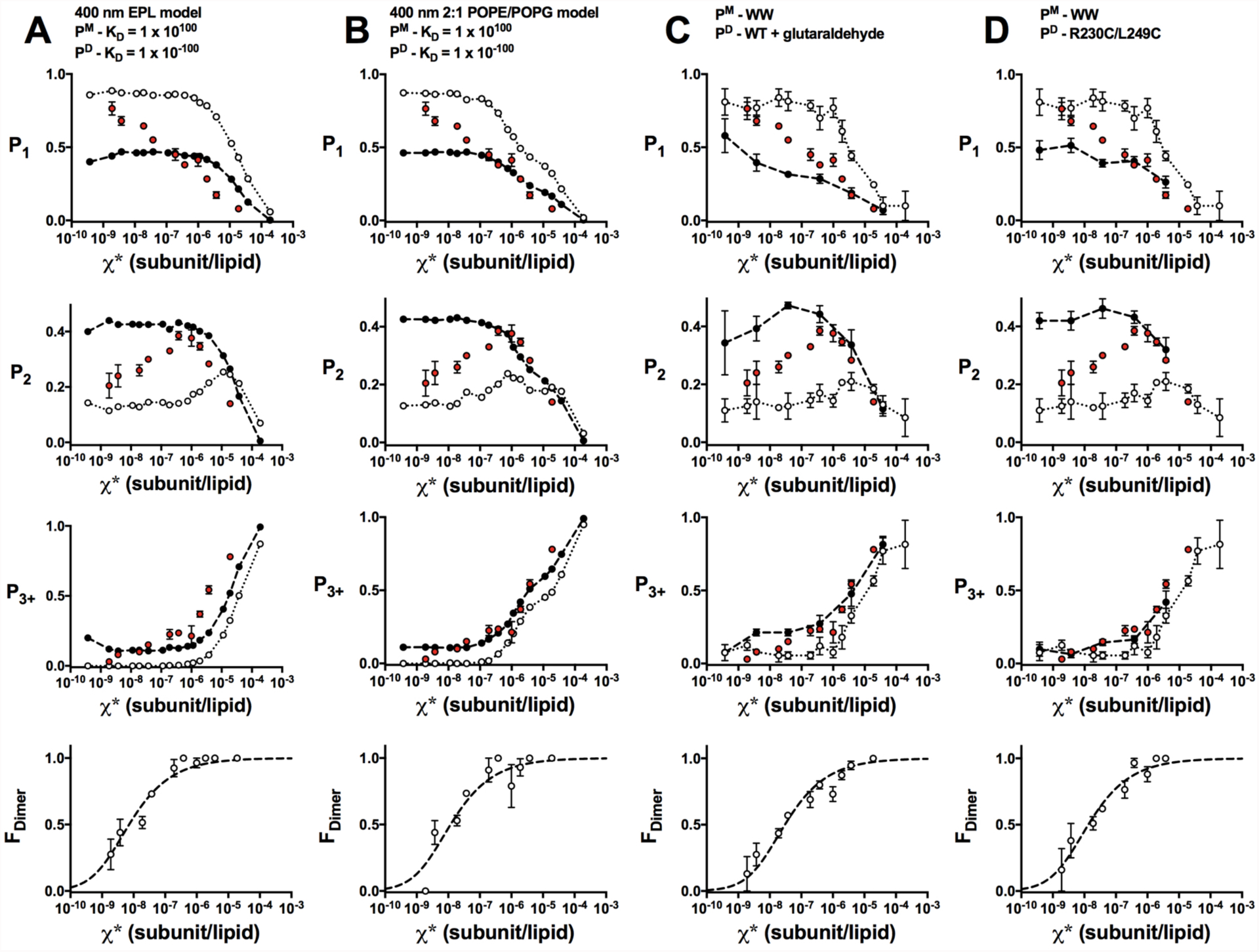
F_Dimer_ analysis of WT-Cy5 in 2:1 POPE/POPG membranes. Photobleaching probabilities (P_1_, P_2_, P_3+_) and F_Dimer_ analysis (bottom row) of WT-Cy5 (red) along with monomer (white) and dimer (black) probabilities. Monomer and dimer probabilities determined by (A) modeling using the 400 nm EPL Walden liposome size distribution (model parameters set as: P_Cy5_ = 0.72, P_non-specific_ = 0.14, bias = 4 and A_lipid=0.6_), ΔG° = -11.1 ± 0.1 kcal/mole, R^2^ = 0.91 (B) modeling using the 400 nm 2:1 POPE/POPG liposome size distribution, ΔG° = -10.8 ± 0.2 kcal/mole, R^2^=0.80 (C) WW-Cy5 for monomer and glutaraldehyde cross-linked WT-Cy5 for dimer, ΔG° = -10.3 ± 0.1 kcal/mole, R^2^ = 0.88, and (D) WW-Cy5 for monomer and R230C/L249C-Cy5 for dimer, ΔG° = -10.8 ± 0.1 kcal/mole, R^2^ = 0.89. Data represent mean ± SEM.

**Figure 3 - figure supplement 2.**
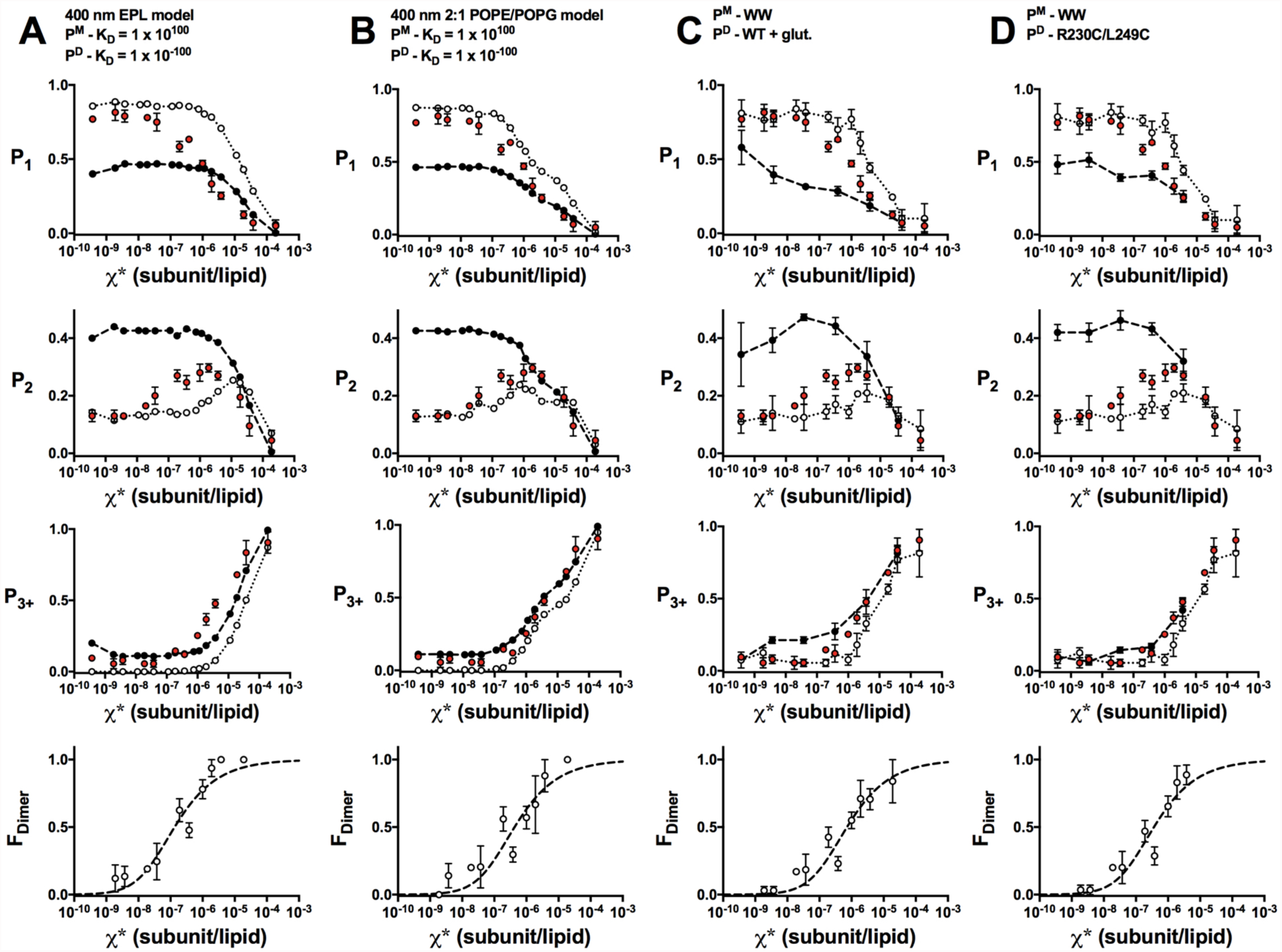
F_Dimer_ analysis of W-Cy5 in 2:1 POPE/POPG membranes. Photobleaching probabilities (P_1_, P_2_, P_3+_) and FDimer analysis (bottom row) of W-Cy5 (red) along with monomer (white) and dimer (black) probabilities. Monomer and dimer probabilities determined by (A) modeling using the 400 nm EPL Walden liposome size distribution (model parameters set as: P_Cy5_ = 0.72, P_non-specific_ = 0.14, bias = 4 and A_lipid_=0.6), ΔG° = -9.4 ± 0.2 kcal/mole, R^2^ = 0.87 (B) modeling using the 400 nm 2:1 POPE/POPG liposome size distribution, ΔG° = -8.7 ± 0.2 kcal/mole, R^2^=0.71 (C) WW-Cy5 for monomer and glutaraldehyde cross-linked WT-Cy5 for dimer, ΔG° = -8.4 ± 0.2 kcal/mole, R^2^ = 0.78, and (D) WW-Cy5 for monomer and R230C/L249C-Cy5 for dimer, ΔG° = -8.8 ± 0.2 kcal/mole, R^2^ = 0.80. Data represent mean ± SEM.

